# Hidden information on protein function in censuses of proteome foldedness

**DOI:** 10.1101/2021.02.24.432609

**Authors:** Dezerae Cox, Ching-Seng Ang, Nadinath B. Nillegoda, Gavin E. Reid, Danny M. Hatters

## Abstract

Methods that assay protein foldedness with proteomics have generated censuses of protein folding stabilities in biological milieu. Surprisingly, different censuses poorly correlate with each other. Here, we show that methods targeting foldedness through monitoring amino acid sidechain reactivity also detect changes in conformation and ligand binding. About one quarter of cysteine or methionine sidechains in proteins in mammalian cell lysate increase in reactivity upon chemical denaturant titration consistent with two-state unfolding. Paradoxically, up to one third decreased reactivity, which were enriched in proteins with functions relating to unfolded protein stress. One protein, chaperone HSPA8, displayed changes arising from ligand and cofactor binding. Unmasking this hidden information should improve efforts to understand both folding and the remodeling of protein function directly in complex biological settings.

**One Sentence Summary:** We show that proteome folding stability censuses are ill-defined because they earmark hidden information on conformation and ligand binding.

## Main text

The maturation of an active protein is typically reliant upon the nascent polypeptide folding into a complex topology. Folding involves a thermodynamic component, which describes the free energy difference between the folded state and unfolded state at equilibrium (folding stability, ∆*G*). Folding also involves other sequential processing steps, such as post-translational modifications and transport. Because folding is fraught with potential mishaps including misfolding and aggregation, in cells a “proteostasis” network oversees all steps related to synthesis, folding and degradation (*1*). In mammalian cells this network consists of several hundred proteins, including molecular chaperone families (e.g. heat shock protein families 40, 70 and 90), the ubiquitin-proteosome system, autophagy and stress response systems (*2*).

Proteostasis imbalance is implicated in diseases involving inappropriate protein aggregation, including neurodegeneration (*3*). As such, there has been extensive interest in determining how protein foldedness varies for proteomes inside cells in healthy and disease contexts (*1, 4, 5*). A canonical approach for measuring protein folding stability of a purified protein involves measuring the abundance of folded and unfolded states in different concentrations of chaotropes, such as urea or guanidine hydrochloride, or exposure to increasing temperature. This approach yields measures of ∆*G* or other correlates of ∆*G* such as chemical denaturation midpoint (*C*_*m*_) or thermal melting midpoint (*T*_*m*_) values that are informative in the case where the protein folds by a two-state equilibrium mechanism. Recent advances have allowed this canonical strategy to measure protein folding stabilities in biological extracts, thereby enabling the *en masse* determination of folding stabilities of proteins (*6*–*13*). Measurement strategies have targeted differences between the folded and unfolded states, such as aggregation propensity, sensitivity of solvent exposed amino acid sidechains to reactive chemicals, which we hereon refer to as residue labeling, and protease cleavage susceptibility (*14*).

Despite all this new information on protein folding censuses, it remains untested how generally applicable the approaches used for studying purified proteins are when applied to complex cellular milieu. In biological settings these methods are also likely to report on other features of proteins including conformational change, ligand binding and protein network organization. We investigated this question and hereon report remarkable additional complexity in the data, as well as a strategy to unmask hidden information pertaining to protein function in complex biological milieu.

### Methodological differences yield poorly correlated measures of thermodynamic proteome stability

First, we examined 20 datasets from 12 studies that reported high-quality protein folding stability data, which used limited proteolysis, residue labelling and thermal profiling methods to assay foldedness (complete reference details are provided in Table S1). These studies reported *T*_*m*_ or *C*_*m*_ values which, for two-state folding models indicate the conditions of equal populations of unfolded and folded protein (i.e. where ∆*G* equals zero). We hypothesized that if these values authentically report on two-state protein folding stability the different datasets should correlate with one another. To test this hypothesis we first scaled each *T*_*m*_ or *C*_*m*_ dataset to range between 0 and 1, to account for the inherently different magnitudes of *T*_*m*_ and *C*_*m*_ values, and then performed pairwise comparisons between datasets. Linear regressions fitted to each comparison revealed a strong positive relationship in stabilities when comparing datasets derived from the same methodology, particularly among thermal profiling data (Fig. 1; lower diagonal datasets 7 - 20). This conclusion was not dependent on the species from which the protein stability was measured, suggesting that closely related proteins from different species behave similarly in terms of the *T*_*m*_ and *C*_*m*_ values. However, comparisons between the datasets derived from different methodologies, such as thermal profiling versus residue labelling, revealed poor correlations at best and none at worst. This lack of correlation was supported by Spearman’s correlation coefficients calculated for each comparison (Fig. 1; upper diagonal), whereby significant positive correlations were primarily observed between datasets derived from the same methodology. One notable exception was residue labeling dataset 4 (*15*), which was moderately correlated with 14 of the 15 thermal profiling datasets. However, closer inspection of dataset 4 revealed that only 17% of the quantified data reported *C*_*m*_ values, because these were the only data that fitted well to a two-state unfolding curve. It thus follows that 83% of the data was disregarded as not fitting to the two-state model (see Table S1 for complete reference details). By contrast, between 66% and 99%, or 66% and 69% of proteins were reported to be well fitted to two-state unfolding isotherms in other thermal profiling and residue labelling datasets, respectively. Collectively, these findings suggested that the most canonical and simple patterns of unfolding, i.e. those that look like two-state unfolding curves were indeed the most likely to report on two-state-like protein unfolding behavior and to correlate with different apparent folding stability datasets. More intriguing however was that such two-state like stability values encompass only a fraction of the data available. Hence the remaining data likely was more applicably explained by complex unfolding mechanisms or mechanisms distinct to folding.

**Fig. 1:**
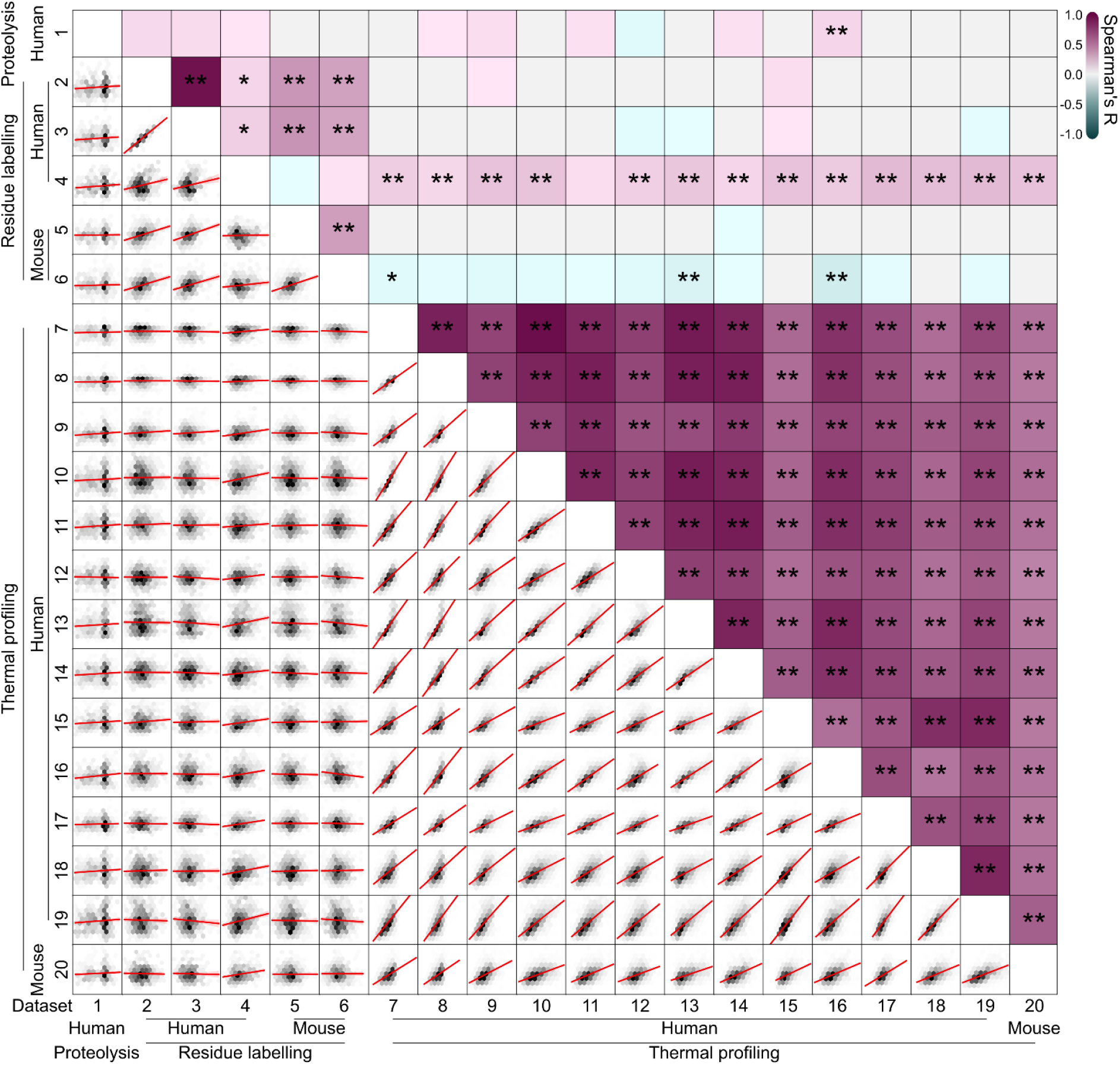
Limited correlations between measures of folding in published proteome stability datasets. Shown are pairwise cross-correlations of normalized protein folding stability measures (*T*_*m*_ or *C*_*m*_). The bottom diagonal shows hexbin density plots, where data point density tiles (grayscale) are overlayed with the fitted linear regression and corresponding 95% C.I. (red; in most cases intervals are too small to be seen). The upper diagonal shows pairwise Spearman’s coefficients (R) represented in the form of a heatmap, overlayed with significance denoted by * (p < 0.05) or ** (p < 0.01). Datasets are ordered according to species of origin (human or mouse) and method of stability measure (limited proteolysis, residue labelling or thermal profiling). Datasets were derived from the following publications; Leuenberger et al., 2017, Science (1), Ogburn et al., 2017, J Proteome Res. (2,3), Walker et al., 2019, PNAS (4), Roberts et al., 2016, J Proteome Res. (5,6), Jarzab et al., 2020, Nat Methods. (7, 8, 9, 16, 17, 20), Becher et al., 2016, Nat Chem Biol. (10), Franken et al., 2015, Nat Protoc. (11), Miettinen et al., 2018, EMBO J. (8), Savitski et al., 2018, Cell. (13, 14), Ball et al., 2020, Commun Biol. (15), Savitski et al., 2014, Science. (18), Sridharan et al., 2019, Nat Commun. (19). Complete reference information is provided in Table S1.

### Residue labelling techniques reveal nuanced and heterogenous changes in protein conformation due to chemical denaturation

To further examine which data can be appropriately explained in terms of two-state folding or not, we collected our own dataset of apparent folding stability using tetraphenylethene maleimide (TPE-MI) as a probe for unfolded proteins. TPE-MI reacts with exposed cysteine free thiols that are otherwise buried in the folded state (*6*). Free cysteine thiols are the least surface-exposed residue of all amino acids in globular proteins so provide an excellent target for examining protein foldedness (*16*). We first performed a urea denaturation curve of purified β-lactoglobulin, which is a model globular protein containing a single buried free thiol residue, and unfolds via two-state-like behavior (Fig. S1). The rate of reaction of TPE-MI with β-lactoglobulin was proportional to the anticipated exposure of the buried thiol upon two-state unfolding. The relationship between rate of reaction and urea concentration yielded a *C*_*m*_ consistent with that obtained from intrinsic tryptophan fluorescence and in accordance with other published results on β-lactoglobulin folding (*17*).

To use TPE-MI on a proteome-wide scale we created denaturation curves of cell lysate with urea (Fig. 2A). Lysates were prepared from mouse neuroblastoma cells (Neuro2a) subjected to Stable Isotope Labeling by Amino acids in Cell culture (SILAC) using light or heavy (^13^C L-lysine and ^13^C,^15^N L-arginine) isotopes for quantitation of reactivity. In essence, lysate from light-isotope labeled cells was used as the “native” control versus lysate from heavy-isotope labeled cells prepared with different concentrations of urea. The light and heavy-isotope labeled samples were each reacted with TPE-MI before mixing and quantitation for the level of reactivity. The extent of cysteine reactivity was determined from the change in abundances of peptides with unreacted cysteines normalized to the ratio of peptides from the same protein that lacked cysteine (Fig. 2A).

**Fig. 2:**
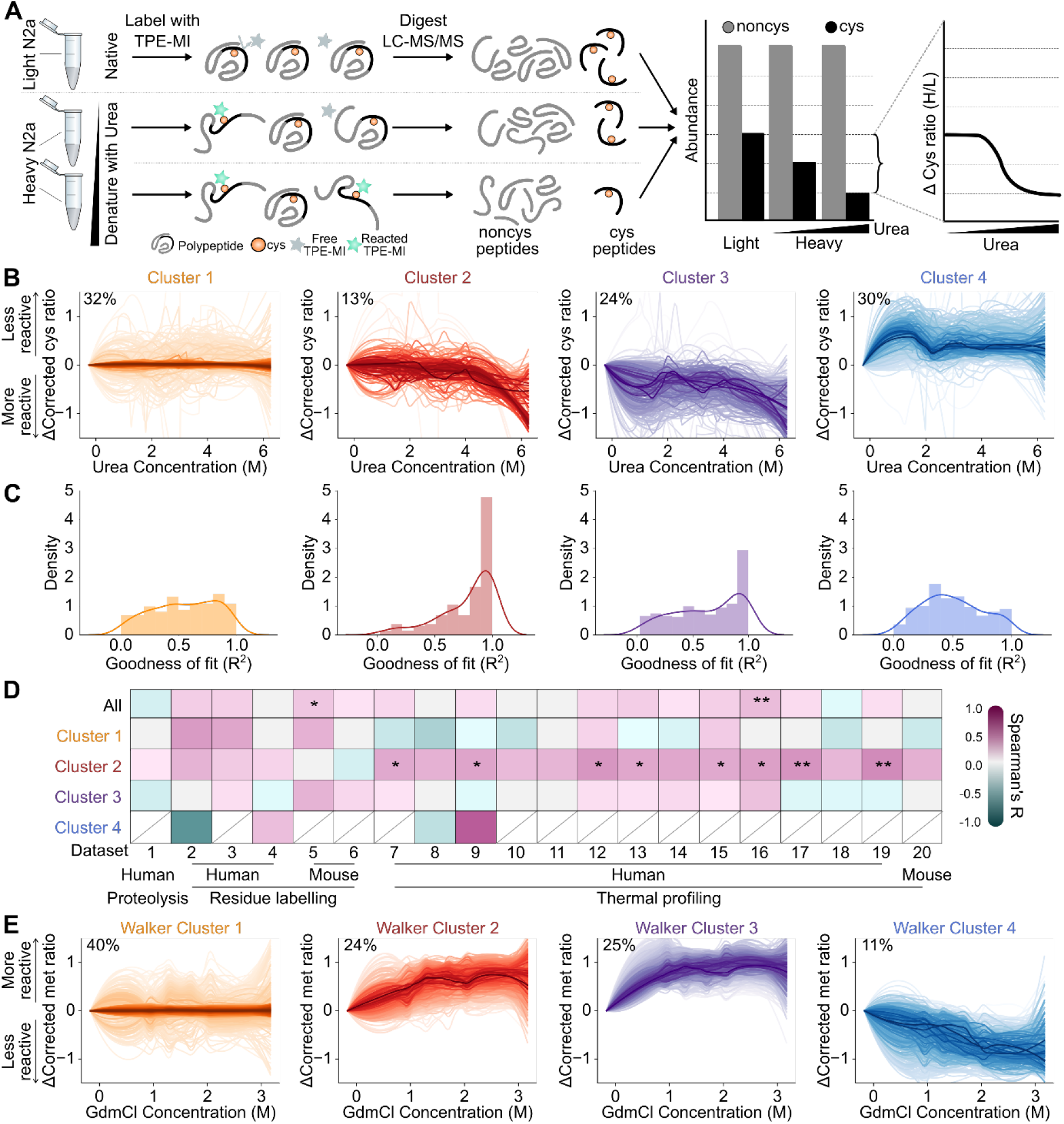
Residue labelling methods reveal patterns of change inconsistent with two-state unfolding during chemical denaturation. (A) Schematic representation of the workflow used to detect conformational change using TPE-MI following chemical denaturation. (B) Clustering of cysteine-peptide TPE-MI reactivity profiles as a function of urea concentration using fuzzy c-means. Families of individual peptide traces are shown where the color saturation reflects the cluster membership score for each peptide, such that the darkest traces are representative of the typical response for each cluster. Data (*n*=3) were smoothed by significance scaling and loess smoothing. The proportion of total peptides assigned to each cluster is also indicated. (C) Histogram of goodness-of-fit among individual peptides following fitting to a two-state unfolding curve. (D) Cross-correlation analysis of peptides in each cluster to previously published stability datasets (list of datasets in Table S1). Correlation heatmap is colored according to the Spearman’s correlation coefficient (R) and significance is denoted by * (p < 0.05) or ** (p < 0.01). Datasets are ordered according to species of origin (human or mouse) and method of stability measure (limited proteolysis, residue labelling or thermal profiling). Missing points (gray line) indicate fewer than 5 proteins in common. (E) Corresponding clustering analysis of an independent published residue labeling dataset (dataset 4) that targeted oxidation of exposed Met residues (15). Data here is represented as per panel B.

To determine the underlying trends in cysteine reactivity as a function of urea concentration we clustered the cysteine peptide reactivity profiles using an unbiased computational approach of fuzzy-c means (*18, 19*). This analysis yielded four distinct patterns of cysteine response to urea titration (clusters 1–4) shown in Fig. 2B. Cluster 1 was defined by no systematic change in reactivity with urea concentration. Clusters 2 and 3 both showed a progressive increase in cysteine thiol reactivity, which was anticipated for greater exposure of buried thiols upon denaturation. Cluster 2 differed from cluster 3 by showing reactivity changes first occurring at higher urea concentrations whereas the changes occurred at lower urea concentrations for cluster 3. Cluster 4, representing one third of the identified cysteine peptides, revealed a systematic decrease in reactivity upon increasing concentrations of urea. This was counter to the anticipated increase in reactivity expected from the exposure of buried cysteine residues induced by denaturation, suggesting that distinct processes were occurring in some proteins in response to urea titration that led to greater burial of exposed cysteine thiols (discussed in more detail below).

To determine which of the data were most consistent with a two-state folding mechanism, individual cysteine reactivity curves were fitted to a two-state unfolding curve and assessed for goodness of fit (Fig. 2C). Consistent with the greater reactivity upon urea denaturation, clusters 2 and 3 contained the most peptides with good fits defined by an R^2^ > 0.9 (Fig. 2C). However, the peptides in cluster 2 were the only group that showed a significant correlation between the fitted *C*_*m*_ values with those of existing census datasets described herein previously in Fig. 1 (Fig. 2D). Given that the published datasets were pre-filtered in the original studies to be consistent with two-state unfolding, this finding demonstrates an authenticity in this subset of data for tracking bona fide two-state unfolding events. The peptides in cluster 3 did not share this correlation with other datasets, and may be a diagnostic of reactivity reporting on non-two-state folding mechanisms such as multistate or non-reversible folding, or other non-folding related mechanisms (Fig. 2D).

To investigate whether the general conclusions made from the TPE-MI dataset can be drawn in datasets derived from other residue labeling methods for foldedness, we re-examined the pre-processed peptide quantitation from dataset 4 (*15*) according to our clustering procedures. Dataset 4 monitored methionine exposure in the lysate of human cell line, HCA2-hTert, by a free radical oxidation approach in different concentrations of the denaturant guanidine hydrochloride (*15*). This dataset was therefore analogous to the TPE-MI approach but independent in multiple parameters of chemical denaturant (guanidine hydrochloride versus urea), species (human versus mouse), target residue for labelling (methionine versus cysteine) and research team (independent research labs).

Despite these differences in parameters, the clustering procedures resulted in a strikingly similar grouping of the methionine oxidation data to the TPE-MI dataset (Fig. 2E – note however the direction of change is inverted due to the nature of the measurements) that illustrates several fundamental consistent conclusions. First was that the data formed 4 clusters with similar patterns of response.

Second was that there were broadly consistent proportions of peptides in the different clusters. Most notably was the consistent cluster for increased protection upon denaturant titration that indicated changes inconsistent with unfolding.

Together the TPE-MI and methionine oxidation data indicated that with an unbiased clustering procedure only about one quarter of residue labeling data in chemical denaturation curves of mammalian proteomes can be confidently described as consistent with a two-state unfolding mechanism. Most notably, this independent dataset confirmed the observation that a substantial portion of the reactivity changes measured in the presence of denaturant cannot be satisfactorily explained by unfolding.

### Residue labeling patterns earmark protein conformational rearrangement as well as unfolding

To investigate protein properties that explain the different patterns of response to urea titration, we first examined the physicochemical characteristics of the peptides in each cluster (and the proteins to which they belong to; Fig. 3A). We examined properties predicted from amino-acid composition that pertained to likely burial of the target residues in the folded state, including charge, hydrophobicity, likelihood to reside in regions of secondary structure and relative solvent exposure for individual peptides associated with each cluster. Several characteristics stood out. Notably the peptides in clusters 2 and 3, which we have thus far demonstrated to have best consistency with two-state folding mechanisms, were most likely to contain hydrophobic and solvent-buried residues, and least likely to be in unstructured protein regions which is in accordance with this conclusion (Fig. 3B). For the parent proteins that have known structures (around 20% of the proteins identified), the extent of solvent exposure of the labelled cysteine residue further supported the conclusion that cysteines in cluster 2 and 3 were in buried regions of proteins and hence became exposed upon denaturation (Fig. S2A). For completeness of analysis, we did not identify any enrichment for other protein features, such as whether the peptides resided in active sites, binding sites, functional motifs, disulfide bonds or annotated domains (Fig. S2B–D). While the proportion of residues located within annotated PFAM domains was relatively high (more than 70% in every cluster), this was anticipated due to the likelihood of free thiols being buried in the folded state of most proteins (*20*).

**Fig. 3:**
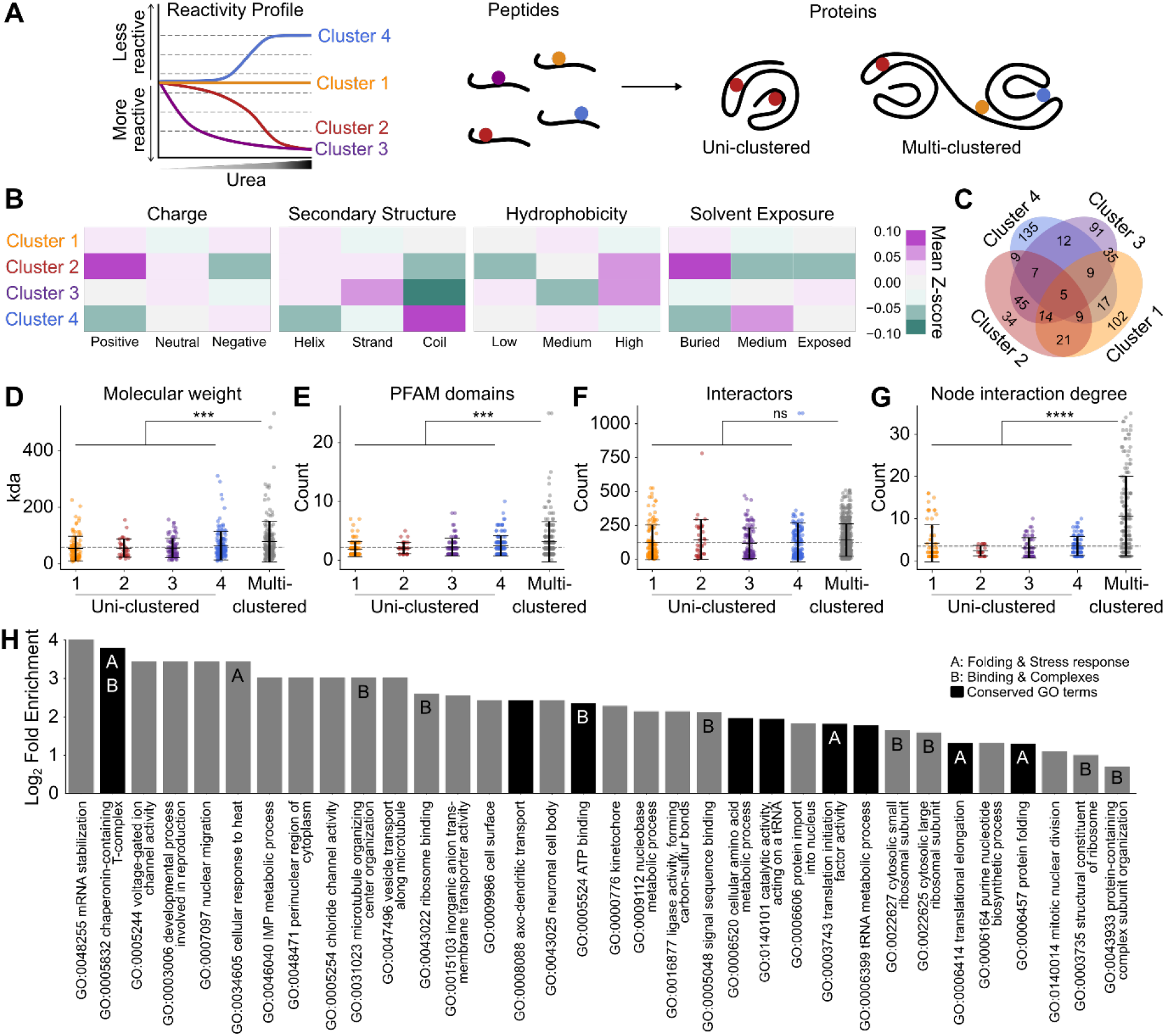
Functional responses to denaturation drive heterogeneous changes in reactivity within single proteins. (A) Schematic overview of peptide cluster patterns and how proteins are grouped depending on the composition of cysteine peptides from multiple clusters. The left graph shows a schematic reactivity profile of each of the four clusters. Peptides are then assigned to their parent protein, which may be deemed “uni-clustered” if only consisting of cysteine-containing peptides from a single cluster, or “multi-clustered” where cysteine-containing peptides from a single protein are associated with more than one cluster. (B) Mean z-score for predicted/extracted physiochemical features according to peptide amino acid composition. (C) Venn diagram depicting proportion of proteins for which peptides were found in each cluster combination. (D) Molecular weight of and (E) number of annotated PFAM domains in proteins to which clustered peptides are assigned. (F) Number of high-confidence first-shell protein-protein interactions and (G) inter-cluster node interaction degree annotated in STRINGdb (v11.0, score > 0.7) for proteins found in each cluster. (H) Gene ontology terms enriched among multi-clustered proteins. Enrichment was determined using Panther GOSlim Fisher’s over-representation test with false-discovery rate correction. Common themes are denoted; A = protein folding and stress response, B = binding and complexes. Dark bars denote exact terms found to also be enriched among multi-clustered proteins in published dataset 4. Panels D-G show individual protein datapoints overlayed with mean ± S.D. Mean of combined uni-clustered proteins is shown as dotted grey line. Uni-clustered proteins were compared to those associated with multiple clusters via t-test with Welch’s correction, *** denotes p < 0.001, **** denotes p < 0.0001, ns denotes p > 0.05.

Because individual proteins may include multiple cysteine-containing peptides that fall into different clusters, we next grouped proteins into categories based on which cluster the peptides belonged to (Fig. 3A). More than half of the peptides belonged to proteins that contained at least one other cysteine peptide from a different cluster (Fig. 3C). Proteins with such peptides were considered “multi-clustered” and separated from proteins that contained cysteine peptides exclusively in one of the other four clusters (which we hereon call “uni-clustered”) for further analysis. Of the multi-clustered proteins, one-third had at least one cysteine peptide which decreased in reactivity (i.e. was in cluster 4) and one or more cysteine peptides for which reactivity increased in the presence of higher concentrations of urea, which suggested multimodal impacts on the protein during denaturation. It is reasonable to predict that such proteins are larger and multi-domain. Indeed, multi-clustered proteins were more likely to have a larger molecular mass and contain more annotated PFAM domains than uni-clustered proteins, consistent with this conclusion (Fig. 3D–E; two-sample t-test, p<0.001). Also of note was the consistency in predicted physicochemical features for uni-clustered proteins whose peptides were associated with cluster 2, which featured elevated hydrophobicity and lower solvent exposure compared to the other uni-clustered and multi-clustered categories (Fig. S3E). This result further supported the conclusion that proteins containing solely cluster 2 peptides were the most likely to display two-state unfolding and that the other proteins with mixed clusters displayed more complex unfolding or other non-folding changes in response to urea titration.

In addition to physicochemical properties, we also examined the molecular functions of proteins assigned to the uni- and multi-clustered protein categories. Protein-protein interaction analysis using the STRING database revealed no difference in the number of direct high-confidence protein binding partners between uni-versus multi-clustered proteins (Fig. 3F). However, the average node degree (the number of protein-protein interactions within each cluster) was up to four-fold higher in multi-clustered proteins than among uni-clustered proteins (Fig. 3G). This result was consistent with the anticipation that multi-clustered proteins are more likely to be multi-domained and larger in size. It therefore follows that such proteins would operate in larger functional networks, which display coordinated changes in cysteine reactivity. By comparison, uni-clustered proteins, particularly those in cluster 2, were more likely to be poorly interconnected. This conclusion is consistent with an anticipation that these proteins were simpler globular proteins whereby the data reported solely on their foldedness and not their function in networks.

Gene ontology (GO) analysis was investigated for each of the protein categories to examine the possibility of a coordinated functional response corresponding to the cysteine reactivity changes (Fig. 3H). Of the 34 significantly enriched top-level GO terms in the multi-clustered proteins, half encompassed mechanisms pertaining to binding and protein complexes or proteostasis response mechanisms such as protein folding machinery. The enrichment of proteostasis mechanisms was striking in that it pointed to the thiol reactivity changes arising in part as functional responses to the stimulus of denaturation by urea. More specifically, three of the GO terms (chaperonin-containing T-complex, GO:0005832; cellular response to heat, GO:0034605; and protein folding, GO:0006457) related to the stimulus of denaturation. In other words, changes in thiol reactivity appeared to earmark changes in ligand binding or the conformation of select proteins that have functions in responding to unfolded proteins that accumulate at higher concentrations of urea. A parallel analysis of the methionine oxidation data (dataset 4) yielded nine identical GO terms (Fig. 3G for common terms, Fig. S3 for all dataset 4 specific terms). This result was further striking in that data from a distinct method, species and denaturant led to a conserved GO enrichment of terms related to responses to unfolded proteins, including identical terms of chaperonin-containing T-complex (GO:0005832) and protein folding (GO:0006457). These data therefore led us to conclude that multi-clustered proteins were likely to encompass functional responses to protein denaturation. By contrast, GO analysis of uni-clustered proteins showed no conserved terms between the two datasets (Fig. S4).

### Residue reactivity captures features of chaperone conformational change in lysate

To decipher the molecular mechanisms that underlie the functional responses to unfolded protein, we next focused on a class of proteins that we expected to have functions in engaging with unfolded proteins, i.e. molecular chaperones. 42 proteins annotated with the ontology term “chaperone-mediated protein folding” machinery (GO:0061077) were identified across the protein groups, particularly in the multi-clustered proteins (Fig. S5). One of these proteins HSPA8 (HSC70; P63017) is the cognate heat shock protein 70 (Hsp70), which showed 3 cysteine peptides in different clusters (Fig 4A). HSPA8 binds to unfolded proteins in concert with J-domain protein co-chaperones and nucleotide exchange factors (*14*). Together they drive protein folding through ATP-dependent cyclical binding and release (*21*). A conserved structural feature of Hsp70 proteins are four modules: an N-terminal nucleotide binding domain (NBD), a substrate binding domain (SBDβ), a helical lid domain (SBDα), and a disordered C-terminal tail of variable length (*22*). The disordered tail of HSPA8 comprises an EEVD motif that mediates interactions with cofactors such as J-domain protein DNAJB1.

**Fig. 4:**
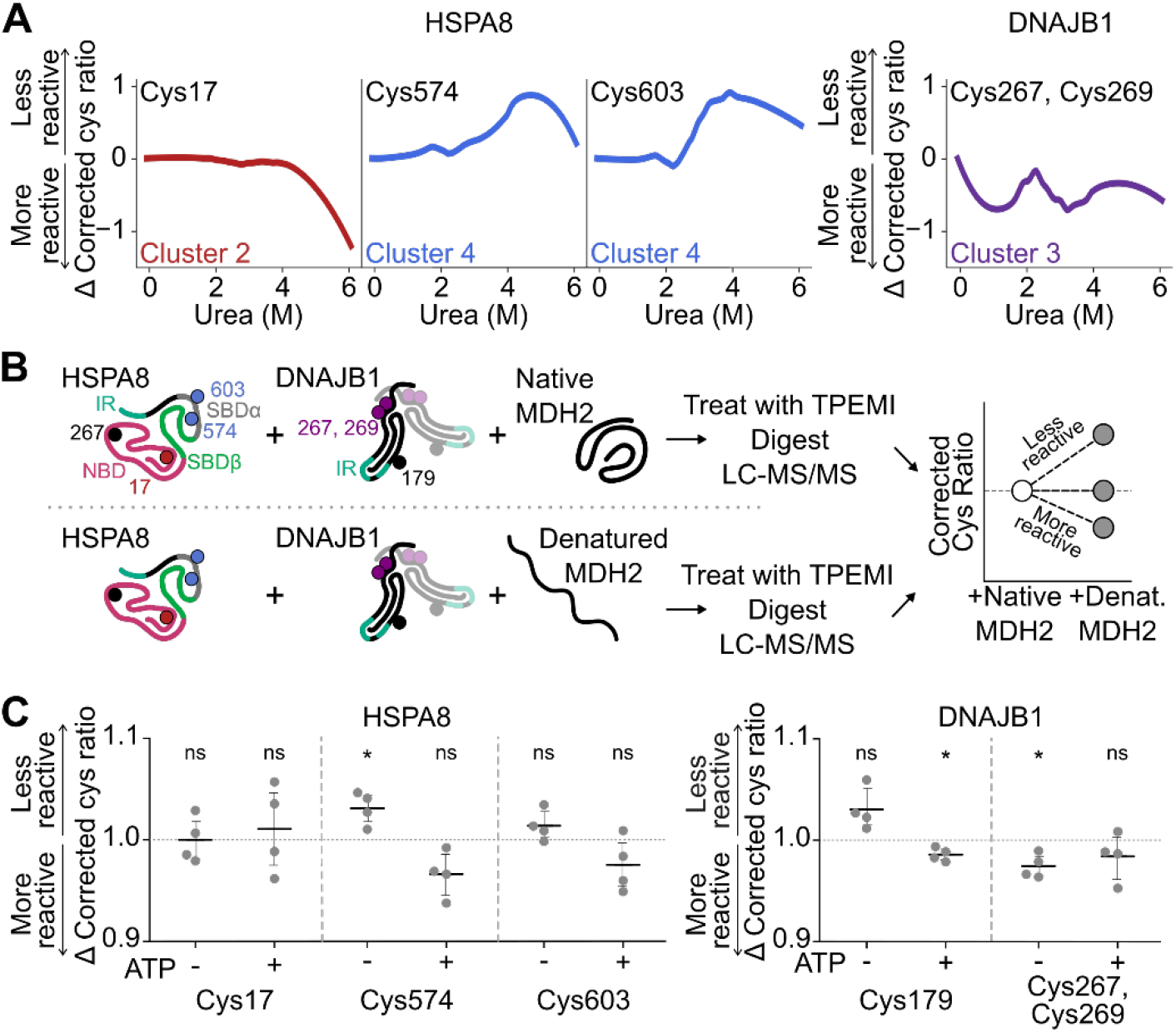
Detection of distinct HSPA8 conformations by residue labeling approaches. (A) Changes in thiol reactivity of HSPA8 and DNAJB1 peptides in Neuro2a lysate titrated with urea, colored according to their assigned cluster. (B) Schematic for recombinant client-binding assay. Human HSPA8 and DNAJB1 were incubated with native or heat-denatured client MDH2. Nucleotide binding domain (NBD; ruby), substrate binding domains (SBDβ; green and SBDα; grey) and cofactor interaction region (IR; teal) are shown on protein backbones, and representative cysteine residues are colored according to the cluster their respective peptides were assigned (orange, red, purple and blue correspond to clusters 1 – 4 respectively, black was not observed). In the case of DNAJB1, dimer is shown with second monomer desaturated. Detailed structural models are shown in Fig. S6. (C) Change in cysteine reactivity of peptides derived from human HSPA8 and DNAJB1 when incubated with heat-denatured MDH2. Recombinant reaction components were incubated in the absence or presence of exogenous ATP prior to TPE-MI labelling. Shown is mean ± S.D. (*n=4*), and deviations from the expected mean of 1 were tested using a one-sample t-test (* denotes p < 0.05, ns denotes p > 0.05).

One peptide from the NBD, containing Cys17, became more reactive to TPE-MI at urea concentrations greater than 4 M (Fig. 4A). The other two peptides were from the SBDα domain, containing Cys574 and Cys603, and these both became more protected in concentrations of urea between 3 – 5 M (Fig. 4A). We also identified one peptide from DNAJB1, that contained two cysteine residues (Cys267, Cys269). These cysteines displayed a complex biphasic (but overall decreased) reactivity profile upon exposure to increasing concentrations of urea (Fig. 4A). We therefore postulated that the changes in cysteine reactivity in HSPA8 and possibly cofactor DNAJB1 upon urea titration may arise due to allosteric conformational changes resulting from their binding and engagement with unfolded proteins. To test this hypothesis, we examined thiol reactivity changes in a reconstituted HSPA8 system *in vitro* that comprised of purified human HSPA8, DNAJB1 and a well characterized model client, malate dehydrogenase (MDH2) (*23*) (Fig. 4B, detailed structural models shown in Fig. S6). To look specifically for changes resulting from interaction with denatured client, TPE-MI reactivity was compared between reconstituted systems containing native versus thermally-denatured MDH2 using tandem mass tag (TMT) isotopic labelling (Fig. 4B).

First, we assessed the system without added ATP. Under this condition HSPA8 and DNAJB1 can bind to non-native MDH2 substrate to form a stable complex (*24, 25*). More specifically, the SBDα domain of an Hsp70 protein (DnaK) has been shown to interact with substrate when the chaperone is in the ADP-bound state (*26*). We saw no change in reactivity of the HSPA8 NBD peptide containing Cys17 (Fig. 4C) suggesting that the increase in reactivity observed above 4 M urea titration in lysate was attributable to NBD unfolding. In contrast, the SBDα domain peptide containing Cys574 decreased in reactivity (one-sample t-test, p=0.033). This result therefore suggested that the decrease in reactivity observed in the SBDα region during urea denaturation arose from substrate and/or ligand binding.

In DNAJB1, we observed two cysteine peptides in the reconstituted system. One peptide, containing Cys179, is located close to the hinge between two β-barrel-like subdomains (CTDI and CTDII) that binds to substrates (Fig. S6). The second, containing two closely-adjacent cysteines (Cys267, Cys269), is located in the homodimer interface. It is important to note that we could not ascribe reactivity changes to a single cysteine within this peptide, thus the changes represent an average across these residues. The peptide containing Cys267 and Cys269 became significantly more reactive under these conditions (Fig. 4E; one-sample t-test, p=0.023). The central location of this peptide within the homodimeric interface of DNAJB1 suggested that accumulated client binding altered the conformation of DNAJB1 to expose the structure nearby these cysteines.

Next we examined the effect of adding supplemental ATP to the reconstituted system, which we predicted would fuel HSPA8 to undergo the full catalytic cycle and therefore release accumulated complexes of HSPA8 and DNAJB1 bound to client (*22*). The Cys574 peptide from HSPA8 became more reactive under these conditions, which is in agreement with HSPA8 disengaging client and/or DNAJB1. Intriguingly, the Cys179 peptide from DNAJB1 showed increased reactivity (Fig. 4C; one-sample t-test, p=0.023). Of note, this peptide is close to the region of DNAJB1 shown to bind the EEVD motif of HSPA8 (*27*), suggesting a level of deprotection when client-bound complexes of the HSPA8-DNAJB1 machinery are dissociated. While this is an attractive hypothesis, we could not exclude the possibility of other allosteric changes associated with DNAJB1 activity. Namely, intramolecular J-domain interactions with the hinge region occur near this residue which are modulated by DNAJB1 engagement with HSPA8 (*23, 28*–*30*). The peptide in DNAJB1 containing (Cys267, Cys269) was no longer protected (Fig. 4C; one-sample t-test, p=0.26), which is consistent with the lack of accumulated substrate-chaperone complexes that would otherwise drive the conformational changes at the dimer interface described above. These results exemplify how residue exposure methodology can track functionally relevant conformational changes in chaperone machinery distinct from unfolding events in complex cellular milieu.

### Conclusions

Collectively, our findings reveal hidden complexity in proteome-wide datasets targeting foldedness with residue labeling approaches. Namely, we demonstrate that a substantial component of the changes seen in residue labeling datasets applied to study proteome denaturation by chemical denaturants are better explained by changes in protein conformation and ligand interactions than unfolding. These findings have two implications of note. First is that analysis of the peptides (and proteins) that generate denaturation curves most similar to two-state unfolding curves provides the most consistent correlation between methodological approaches and provide a more robust core census list of stability measurements. This is a critical point with respect to the retrospective consideration of whole proteome stability datasets because it appears that upwards of two thirds of the data previously fitted to a measure of folding stability may be more appropriately explained by changes in conformation or ligand association. Other proteomics approaches have drawn general conclusions that are in agreement with our findings here, namely how changes in proteome solubility encode information on rewired protein interaction networks (*31*–*33*). Others have also shown that ligand interactions can modulate the thermal melting profiles of proteins in lysate (*8, 34, 35*), and recent commentary speculates this is one among a range of biophysical effects that could contribute to a protein’s observed thermal stability (*36*). The workflow presented here provides a useful strategy to delineate changes arising from unfolding from these other attributes of proteins.

The second implication is that the ability to determine subtle changes in tertiary and quaternary conformation with domain-level resolution distinguishes amino-acid specific methods from thermal melt and aggregation-based techniques. Subtle changes in protein conformation mediate protein-protein interactions underlying many cellular functions. However, the dynamic and often transient nature of these interactions can make them challenging to quantify in cells *en masse* and requires sensitive but non-specific conformational probes capable of distinguishing domain-specific changes. Overall, the data presented here support the ability of residue labelling methodologies such as TPE-MI to fill this void, providing quantitative insight into aspects of protein conformation beyond stability and unfolding. We anticipate this to open the door for studies on proteome structure and function in natural intact biological settings, including live cells or other complex biological milieu.

## Acknowledgements

We thank Professors Paul Gooley and Heath Ecroyd for helpful discussions and careful reading of the manuscript. We also thank Dr. Yuning Hong (La Trobe University) for providing TPE-MI, and the Bio21 Melbourne Mass Spectrometry and Proteomics facility.

## Author contributions

D.C., N.B.N., G.E.R., and D.M.H. designed the research; D.C. and C.A. performed the research and analyzed the data; and D.C. and D.M.H. wrote the manuscript;

## Data and materials availability

The mass spectrometry raw data files and preprocessed identification datasets have been deposited to the ProteomeXchange Consortium via the PRIDE partner repository with the data set identifiers PXD022587 and PXD022640. All other relevant data and analysis code are available from 10.5281/zenodo.4280621 and 10.5281/zenodo.4287767 respectively.

## Supplementary Materials

Materials and Methods

Figures S1–S6

Table S1

External Datasets S2-S4

## Figures

## Supplementary materials for

### This PDF file includes

Materials and Methods

Figure S1: TPE-MI reports on unfolding of recombinant β-lactoglobulin. Relates to Fig. 2.

Figure S2: Residue and protein physicochemical properties. Relates to Fig. 3.

Figure S3: Gene ontology terms enriched among multi-cluster proteins identified in published residue labelling dataset 4 (*15*). Relates to Fig. 3.

Figure S4: Gene ontology terms enriched among single cluster proteins. Relates to Fig. 3.

Figure S5: Enrichment of chaperone machinery among multi-clustered proteins. Relates to Fig. 4.

Figure S6: Structural models for chaperone client-binding reaction components. Relates to Fig. 4.

Table S1: Published proteome stability dataset details

### Other supplementary materials for this manuscript include the following

Dataset S1: Statistical analyses summary *(separate file)*

Dataset S2: Supplementary dataset – lysate denaturation preprocessed peptide data *(separate file)*

Dataset S3: Supplementary dataset – recombinant client-binding assay preprocessed peptide data *(separate file)*

## Materials and Methods

### Materials

All materials were purchased from Sigma-Aldrich (St. Louis, MO, USA) unless otherwise indicated. The mouse neuroblastoma cell line Neuro2a (N2a) was obtained from lab cultures originating from the American Type Culture Collection and screened for mycoplasma contamination. TPE-MI was stored as stocks (10 mM in DMSO) in the dark at 4 °C before use. Recombinant human HSPA8 and DNAJB1 were purified as previously described (*23*).

### Correlation of published proteome stability datasets

Published basal proteome stability datasets were collected as follows. First 813 articles were collected from PubMed keyword searches performed on July 31^st^ 2020 for “thermal proteome profiling”, “thermal proteome unfolding”, “folding stability proteome”, “limited proteolysis proteome”, “proteome denaturation labelling”, “proteome unfolding label” and “SPROX proteome”. Abstracts were filtered manually for evidence of containing primary experimental datasets for protein stability under control conditions derived from either human or mouse samples. Of these 12 papers were selected as suitable. Datasets were then assigned into one of three categories based on methodology: limited proteolysis, residue labelling and thermal profiling. Complete details for the selected resources, including specific supplementary materials files for each dataset, are provided in Table S1.

Datasets were collected from the relevant supplementary materials analyzed with custom scripts written in Python programming language. The logic of the scripts was to collect the reported protein stabilities provided by each source, and where necessary map the protein identifiers to UniProt Accession numbers. Proteins were mapped to KEGG Orthology (KO) identifiers using previously established protocols (*37*) available via cross-referencing from UniProt https://www.uniprot.org/. Stability measures of different datasets were filtered as per the goodness-of-fit criteria used in the original study, then normalized to 1 to enable cross correlation (i.e. to account for different scales for thermal denaturation values (*T*_*m*_) and chemical denaturation values (*C*_*m*_)). Spearman’s correlation coefficients and p-values were calculated in a pairwise manner for all proteins found to be commonly quantified in a given pair of datasets.

### Recombinant β-lactoglobulin denaturation

A stock solution of recombinant β-lactoglobulin was prepared in PBS (pH 7.4), before being diluted to a final concentration of 250 µM in triplicate in urea prepared at concentrations ranging from 0 – 6 M. Samples were then equilibrated in denaturant for 4 h at 25 °C, before being labelled with TPE-MI (or the vehicle control DMSO) at a final concentration of 50 µM. Immediately after addition of the labelling reagent, samples were transferred to clear-bottom 96-well UVStar plate (Grenier BioOne). TPE-MI (350/20 nm ex, 465/20 nm em) and intrinsic tryptophan (295/10 nm ex, 360/20 em) fluorescence was read every 60 seconds for 60 minutes using a CLARIOstar (BMG Labtech) with shaking at 200 rpm for 5 seconds prior to each cycle. In the case of TPE-MI, the first 9 minutes of linear increase in fluorescence were fitted with linear regression to derive the rate of reaction which was used for subsequent fitting. Both the TPE-MI rate and tryptophan fluorescence data was fitted via non-linear least squares regression to a two-state unfolding curve.

### Cell culture

Neuro2a cells were cultured in Dulbecco’s modified Eagle’s medium (DMEM; ThermoFisher Scientific) supplemented with 10% (v/v) fetal bovine serum (ThermoFisher Scientific) and 1 mM L-glutamine (ThermoFisher Scientific). In the case of isotopically labelled cultures (SILAC), cells were cultured in DMEM (Silantes) supplemented with either unlabeled (light) or ^13^C L-lysine and ^13^C,^15^N L-arginine, along with 10% (v/v) dialyzed foetal bovine serum (ThermoFisher Scientific) and 1 mM L-glutamine (Silantes). Cells were cultured in isotopically labeled media for at least 8 doublings prior to use. Cells were maintained at 37 °C in a humidified incubator with 5% atmospheric CO_2_ and were reseeded into fresh culture flasks once at 80% confluency following mechanical dissociation. For plating, cell count and viability were automatically determined using a Countess trypan blue assay (ThermoFisher Scientific). Cells were seeded in 6 or 12 well plates (Corning) and grown for at least 18 h before treatment.

### Lysate preparation and chemical denaturation

Following treatment, cells were washed once in PBS before being mechanically harvested in fresh PBS and centrifuged at 150 *g* for 5 min. Cell pellets were then resuspended in lysis buffer (50 mM Tris, pH 8.0, 0.8 % (v/v) IGEPAL CA-630, 1.5 mM MgCl_2_) containing cOmplete Mini, EDTA-free Protease Inhibitor Cocktail (Sigma) and 250 U benzonase (Sigma), then incubated on ice for 30 min. Lysate was then centrifuged at 20,000 *g* for 10 min to pellet cell debris, and the resultant supernatant transferred to a fresh Eppendorf tube. Total protein concentration was then determined using a Pierce BCA protein assay (Thermo Scientific) with bovine-serum albumin as the mass standard. A standard volume of lysate was distributed to aliquots of urea prepared at concentrations ranging from 0 – 6 M in water from an 8 M stock for which the concentration was determined by measuring the refractive index. In the case of SILAC lysate, light and heavy-labelled samples were combined at a 1:1 ratio prior to denaturation. Samples were then equilibrated in denaturant for 4 h at 25 °C. Following denaturation, lysate aliquots were labelled with TPE-MI to a final concentration of 100 µM for 30 min at 25 °C, then immediately transferred to a 5-fold excess (v/v) of ice-cold acetone and stored at −20 °C overnight.

### Sample preparation for mass spectrometry

Samples were pelleted at 20,000 *g* for 30 min at 4 °C. Protein pellets were solubilized in 100 µl of 8 M urea in 50mM triethylammonium bicarbonate (TEAB), and incubated with shaking at 37 °C for 45 min. Proteins were reduced using 10mM tris(2-carboxyethyl)phosphine, pH 8.0, and alkylated with 10mM iodoacetamide for 45 min, before being digested with 2 µg trypsin (ThermoFisher Scientific) overnight with shaking at 37 °C. Peptides were then desalted via solid-phase extraction using an Oasis HLB 1 cc Vac Cartridge (catalogue number 186000383, Waters Corp., USA) that was pre-equilibrated by washing. Samples were collected in fresh tubes and lyophilized (VirTis Freeze Dryer, SP Scientific). The lyophilized peptides were subjected to another round of BCA assay as above so as to normalize for similar loading onto the mass spectrometer. The final concentration of peptides was 0.1 µg/µl in 2% (v/v) ACN containing 0.05% (v/v) trifluoroacetic acid.

### NanoESI-LC-MS/MS

Samples were analyzed by nanoESI-LC-MS/MS using a Orbitrap Fusion Lumos Tribrid mass spectrometer (Thermo Scientific) fitted with a nanoflow reversed-phase-HPLC (Ultimate 3000 RSLC, Dionex). The nano-LC system was equipped with an Acclaim Pepmap nano-trap column (Dionex—C18, 100 Å, 75 µm× 2 cm) and an Acclaim Pepmap RSLC analytical column (Dionex—C18, 100 Å, 75 µm× 50 cm). For each LC-MS/MS experiment, 0.6 µg of the peptide mix was loaded onto the enrichment (trap) column at an isocratic flow of 5 µl min^−1^ of 3% ACN containing 0.1% (v/v) formic acid for 5 min before the enrichment column was switched in-line with the analytical column. The eluents used for the LC were 0.1% (v/v) formic acid (solvent A) and 100% ACN/0.1% formic acid (v/v) (solvent B). The gradient used (300 nl min^−1^) was from 3–22% B in 90 min, 22–40% B in 10 min and 40–80% B in 5 min then maintained for 5 min before re-equilibration for 8 min at 3% B prior to the next analysis. All spectra were acquired in positive ionization mode with full scan MS acquired from *m/z* 400–1500 in the FT mode at a mass resolving power of 120,000 after accumulating to an AGC target value of 5.00e^5^, with a maximum accumulation time of 50 ms. Lockmass of 445.12002 was used. Data-dependent HCD MS/MS of charge states > 1 was performed using a 3 s scan method, at an AGC target value of 1.00e^4^, a maximum accumulation time of 54 ms, a normalized collision energy of 35%, and with spectra acquired at a 7,500 mass resolving power of 15,000. Dynamic exclusion was used for 45 s.

In the case of TMT-labelled samples, data were obtained on an Orbitrap Eclipse Tribrid mass spectrometer using nanoESI-LC parameters as described above. All spectra were acquired in positive mode with full scan mode full scan MS from *m/z* 300–1600 in the FT mode at 120,000 mass resolving power after accumulating to a target value 5.00e^5^ and with maximum accumulation time of 50 ms. Lockmass of 445.12002 was used. A preferred inclusion list containing doubly and triply masses of tryptic peptides belonging to HSPA8 (P11142), DNAJB1 (P25685) and MDH2 (P00346) was created to increase the coverage of identifiable peptides from these proteins. Data-dependent HCD MS/MS of precursors that matches the inclusion list then other charge states > 1 were performed using a 3 s scan method, 0.7m/z isolation width, target value of 5.00e^4^, a maximum accumulation time of 54 ms, a normalized collision energy of 35% and at a 30,000 mass resolving power (with TurboTMT mode) to resolve the low mass TMT reporter mass. Dynamic exclusion was used for 45 s

### Peptide identification

Initial data analysis of raw data generated during this study was carried out using Proteome Discoverer (v2.1; ThermoFisher Scientific) or MaxQuant (v 1.6.3.4) against the Swissprot Mus Musculus database (downloaded 04/07/2016; containing 16,795 entries). Searches were conducted with 20 ppm mass tolerance for MS, and 0.2 Da for MS/MS, with one missed cleavage allowed and match between runs enabled. Variable modifications included methionine oxidation, N-terminal protein acetylation, N-terminal methionine cleavage and SILAC-Lys6, Arg10, while the carbamidomethylcysteine modification was fixed. The false discovery rate maximum was set to 0.005% at the peptide identification level (actual was 0.005 for each replicate) and 1% at the protein identification level. All other parameters were left as default.

### Ratio correction and scaling

Further analysis was performed with custom Python scripts. The logic was as follows. First, the common contaminant protein keratin was removed. Quantified proteins were considered as those identified by at least two unique peptides, one of which contained a cysteine residue, and the average peptide abundance ratio for the non-cysteine-containing peptides was calculated for each protein at each urea concentration. The mean non-cysteine abundance ratio was used to correct the corresponding cysteine-containing peptide(s) for any change in overall protein abundance caused by the treatment, yielding the corrected cysteine ratio. The corrected cysteine ratio was normalized to the native sample (0 M Urea), such that no change resulting from urea denaturation would yield a ratio of 1.

The resultant data was then scaled with a p-value weighted correction. This correction weights the mean corrected cysteine ratio of biological replicates (*n=3*) according to the relative confidence with which it deviates from the expected value (in this case, 1) as per equation 1:

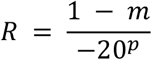

where *m* corresponds to the mean of the corrected cysteine ratios and *p* corresponds to the p-value derived from a one-sample t-test of the corrected cysteine ratios against the expected value at a single denaturant concentration. To improve confidence in the trends of the corrected peptide ratios across urea concentrations, the data were subsequently smoothed with Locally Estimated Scatterplot Smoothing (LOESS). The resultant curves for the peptide ratios across urea concentrations were clustered by using fuzzy-c means, where the optimal number of clusters was first estimated using the *kneed* package before manual inspection of ± 2 centroids to achieve minimal redundancy in clustered patterns. For all subsequent bioinformatic analyses, peptides were then assigned to clusters with the highest membership score.

### Peptide and protein properties

The curves for the peptide ratios across urea concentrations were fitted with a sigmoidal two-state unfolding model as per equation 2:

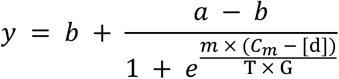

where *a* and *b* correspond to the top and bottom plateaus respectively, *C*_*m*_ corresponds to the denaturant concentration at which both the folded and unfolded states are equally populated at equilibrium (assuming two-state protein folding), *[d]* corresponds to denaturant concentration, and *m* corresponds to the slope. *T* and *G* correspond to the temperature (298.15 K) and gas constants, respectively. Fits were filtered according to the following criteria: (1) R^2^ > 0.75, (2) absolute value of *a* and *b* less than 10, (3) fitted *C*_*m*_ within the range of denaturant concentrations tested, (4) relative error in *C*_*m*_ less than 0.5, and (5) value at 0 M urea in greater than at 6 M urea. The fitted *C*_*m*_ values were then compared against the published datasets as described above.

Physicochemical properties for individual cysteine residues, peptides and proteins of interest were compiled from various databases and extraction/prediction platforms, including UniProt https://www.uniprot.org/, PFAM https://pfam.xfam.org/, Protein Data Bank https://www.ebi.ac.uk/pdbe/, DSSP (*38, 39*), IUPred2A (*40*), STRINGdb (*41*), PantherGOSlim http://pantherdb.org, and iFeature (*42*).

### HSP70 client denaturation assay

Pig heart L-malate dehydrogenase (MDH2; Roche, catalogue # 10127256001) and recombinant human HSPA8 were prepared in HEPES buffer (50 mM HEPES, pH 7.5, 50 mM KCl, 5 mM MgCl_2_, 2 mM DTT) to a final concentration of 5 µM and 2 µM, respectively, in the presence or absence of recombinant human DNAJB1 (1 µM). In the case of heat-denatured samples, MDH2 aliquots were heated to 42 °C for 10 min then returned to 37 °C in a thermocycler (BioRad), while native samples were maintained at 37 °C. MDH2 was then combined with the remaining reaction components in the absence or presence of 2 mM ATP (New England Biosciences, catalogue # P0756S), then incubated at 37 °C for 30 min in a heating block. Samples were then labelled with 100 µM TPE-MI for 15 min at 25 °C before being diluted into 1 ml ice cold acetone and incubated at −20 °C overnight. Samples, including a pooled control sample, were prepared for mass spectrometry using the Preomics iST-NHS (Preomics, catalogue # P.O.00026) and TMT 11-plex labelling (ThermoFisher, catalogue # A37725) kits according to the manufacturer’s protocol. The pooled channel was added to each biological replicate to support efficient normalization between replicates. Resultant peptides were analyzed using a TMT based MS methodology as described above. The collected spectra were searched against a custom database containing sequences for HSPA8 (P11142), DNAJB1 (P25685) and MDH2 (P00346) sequences downloaded from UniProt. The search was conducted as above, with the following alterations: the MS2 reporter ion was set to TMT 11-plex, isotopic distribution correction applied according to the product data sheet and the fixed carbamidomethylcysteine was replaced with the Preomics alkylation (+113.084 Da).

Filtering and further analysis of the dataset was then carried out with custom Python scripts. The logic was as follows. Raw intensities for peptides with no missed cleavages were scaled according to the molar contribution of the corresponding protein to each reaction. The mean peptide abundance ratio for non-cysteine peptides in each protein that were quantified across all channels containing that protein, was then calculated. The non-cysteine intensity was used to correct the corresponding cysteine-containing peptide(s) for any change in overall protein abundance, resulting in the corrected cysteine ratio. In the case of HSPA8 peptides, the corrected cysteine ratio was then normalised to the native HSPA8 sample, and finally the change in corrected cysteine ratio is reported as the mean of two technical replicates.

### Statistical analysis and data availability

Statistical analyses were performed either using the scipy module in python (*43*) or using GraphPad Prism (v 8.4.3). The exact *p* values, raw values and statistical details are provided in Dataset S2.

The mass spectrometry proteomics data have been deposited to the ProteomeXchange Consortium via the PRIDE (*44*) partner repository with the data set identifiers PXD022587 and PXD022640. All other data and analysis code are available from 10.5281/zenodo.4280621 and 10.5281/zenodo.4287767 respectively.

---

**Fig. S1:**
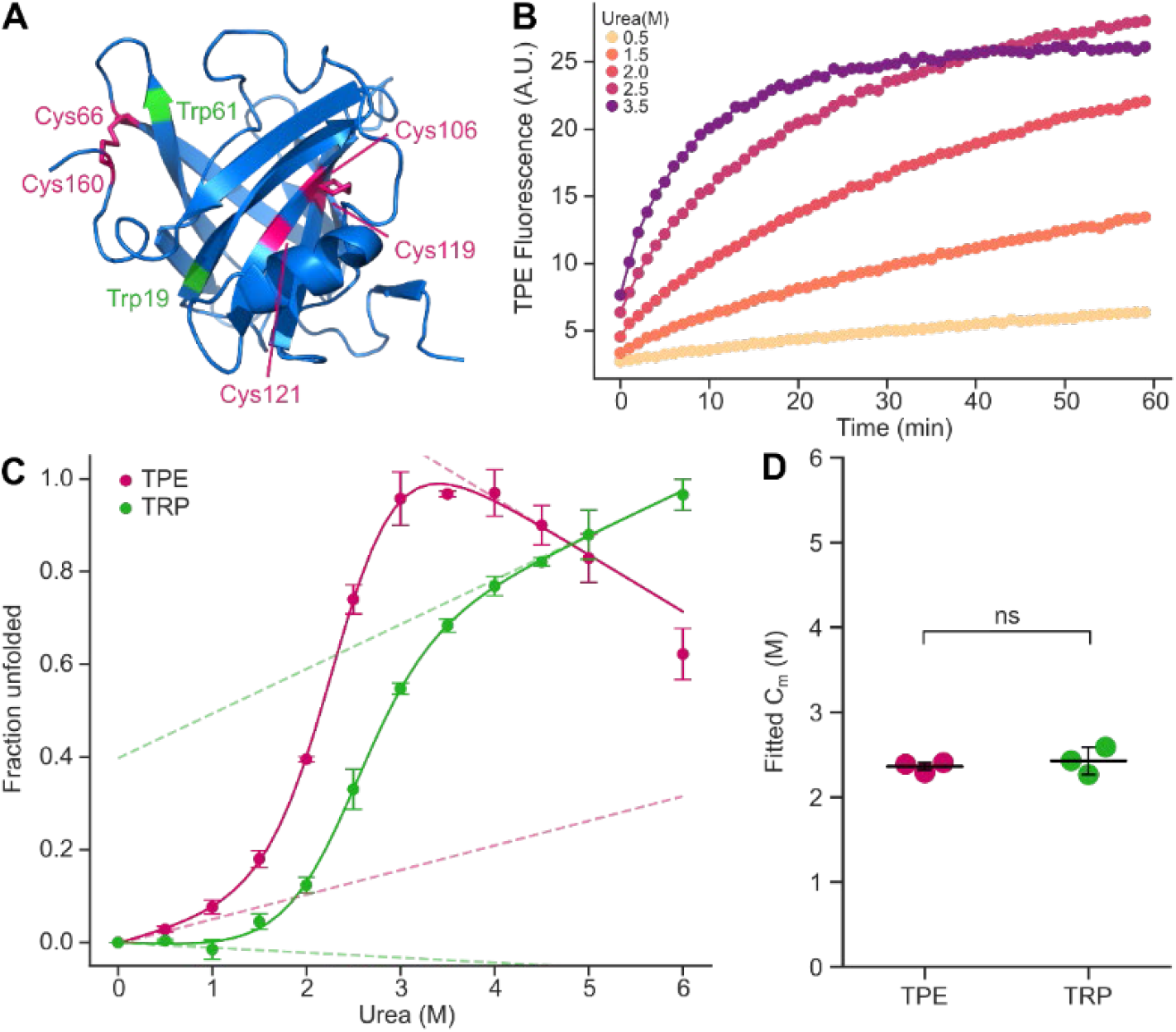
TPE-MI reports on unfolding of recombinant β-lactoglobulin. Relates to Fig. 2. (A) Structure of β-lactoglobulin, a model globular protein, adapted from PDB entry 1CJ5. Pertinent residues are highlighted in magenta (cysteine) and green (tryptophan). (B) Chemical denaturation of recombinant β-lactoglobulin in the presence of TPE-MI. Samples were equilibrated for 4 hours at room temperature before addition of TPE-MI, and TPE-MI fluorescence was monitored every 60 sec for 1 h. (C) The initial rate of reaction is calculated from B via linear regression, and fitted to a denaturation curve. Curve is compared to intrinsic tryptophan fluorescence, also fitted to a denaturation curve. (D) Fitted *C*_*m*_ derived from C, compared using t-test in GraphPad Prism. In panels B – D, data shown is mean ± SD of 3 replicates, and is representative of 2 independent experiments.

**Fig. S2:**
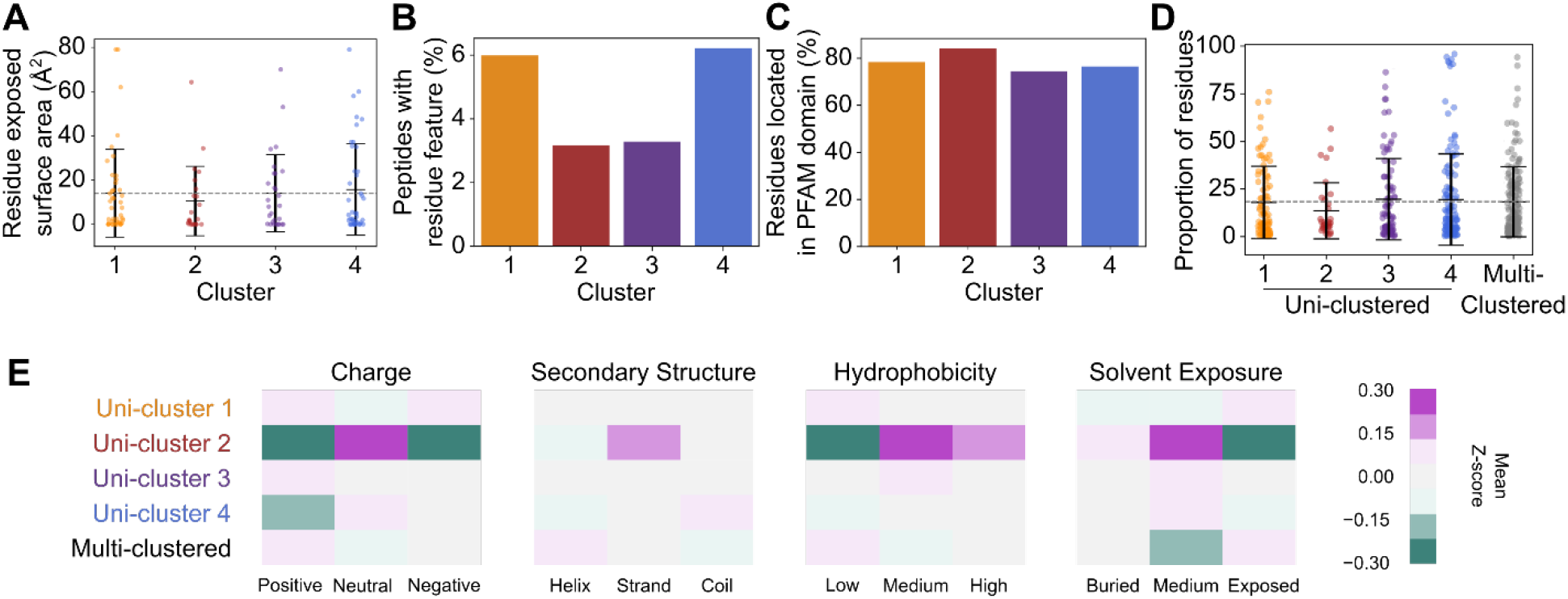
Residue and protein physicochemical properties. Relates to Fig. 3. Individual cysteine residues contained within clustered peptides were assessed for (A) their relative surface exposure in experimentally determined structures available via the Protein Data Bank, (B) the proportion of residues annotated as a functional feature in UniProt, and (C) the proportion of residues located within curated PFAM domains. (D) The proportion of disordered residues in proteins associated with each cluster as predicted by IUPred2. (E) Mean z-score for predicted or extracted physiochemical features according to protein amino acid composition. Panels A and D show individual protein datapoints overlayed with mean ± S.D. Mean of clustered peptides (A) or combined uni-clustered proteins (D) is shown as dotted grey line.

**Fig. S3:**
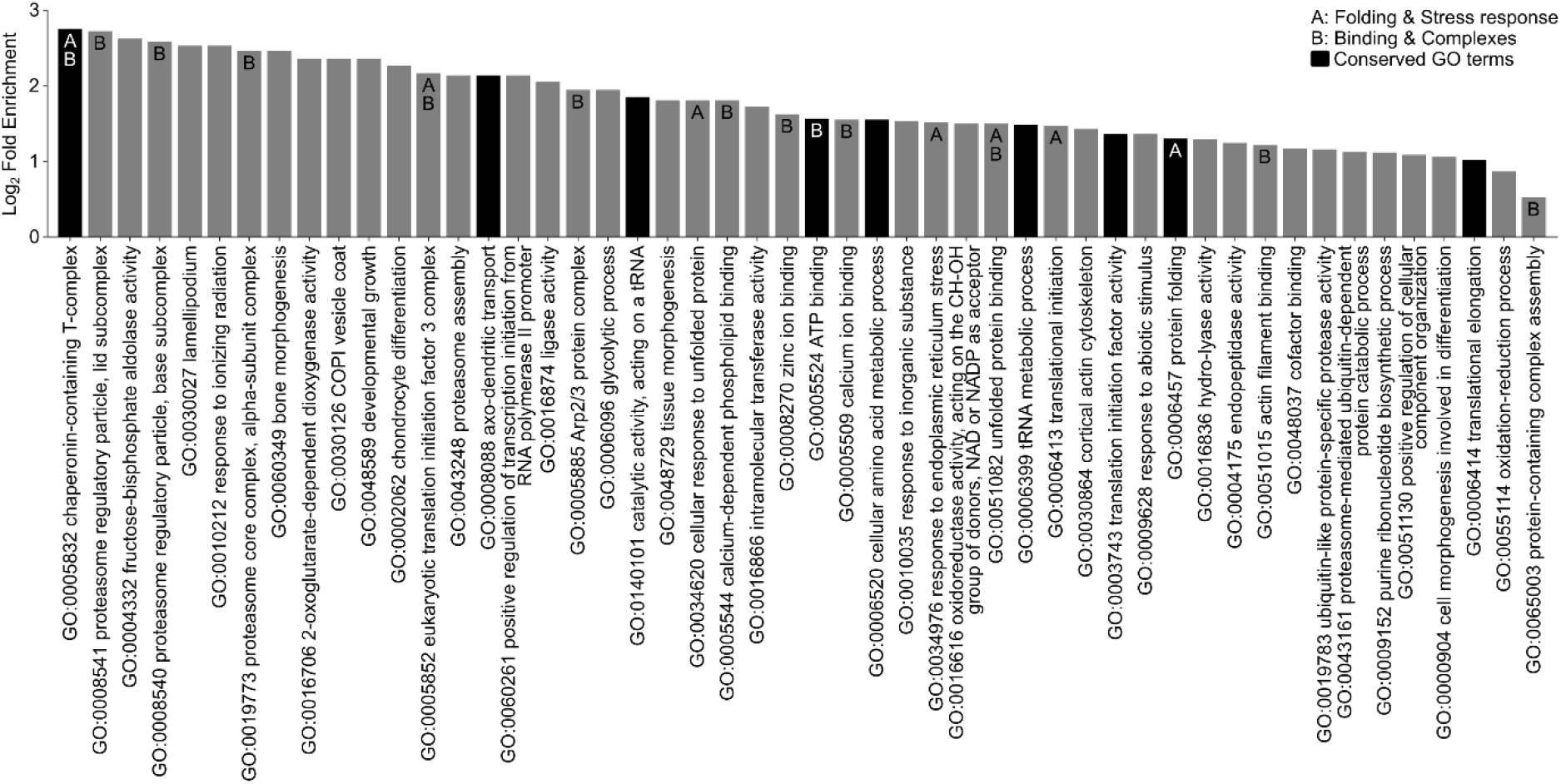
Gene ontology terms enriched among multi-cluster proteins identified in published residue labelling dataset 4 (15). Relates to Fig. 3. Enrichment determined using Panther GOSlim Fisher’s overrepresentation test with false-discovery rate correction. Common themes are denoted; A = protein folding and stress response, B = binding and complexes. Dark bars denote exact terms found to also be enriched among multi-cluster proteins in the TPE-MI dataset.

**Fig. S4:**
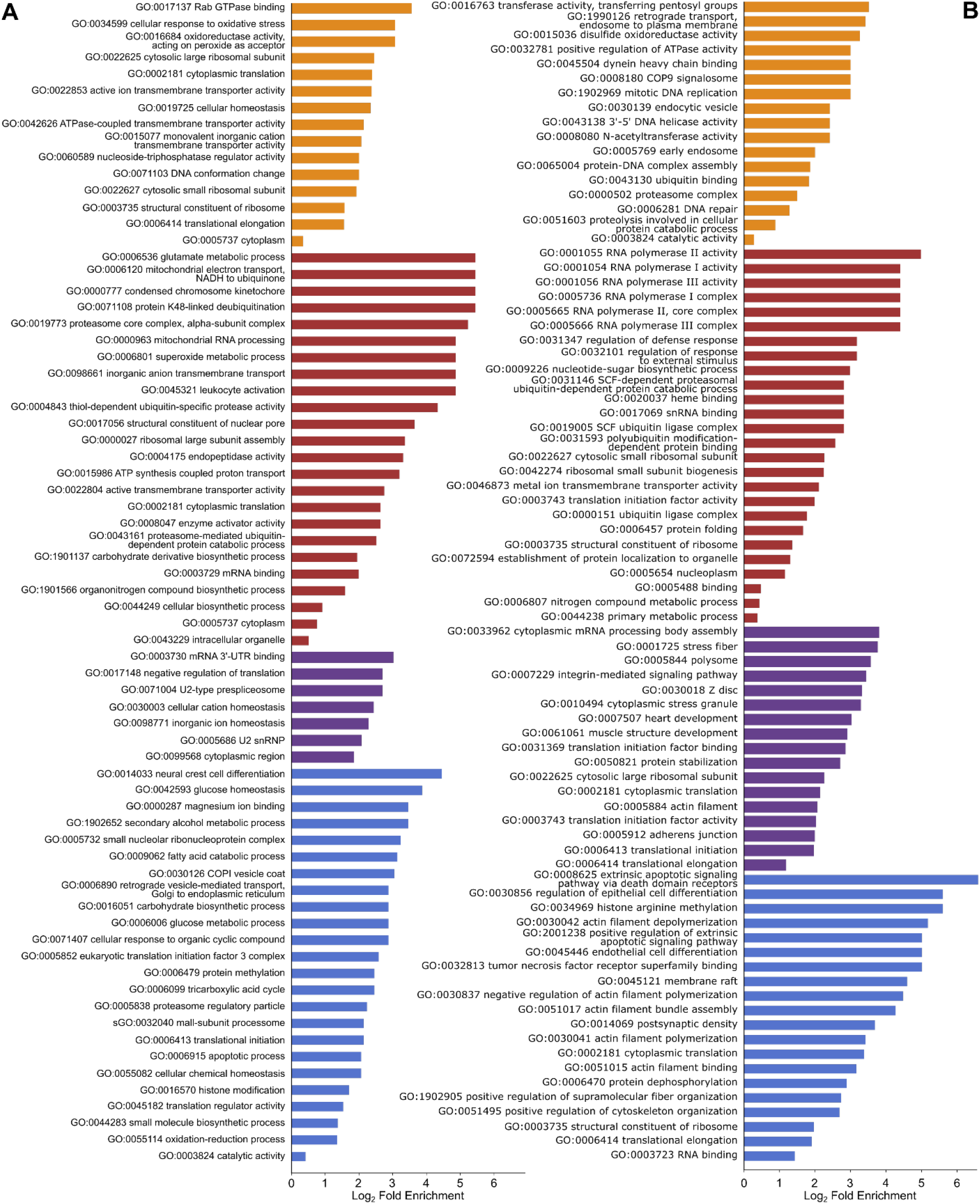
Gene ontology terms enriched among uni-clustered proteins. Relates to Fig. 3. Enrichment for (A) TPE-MI and (B) (15) datasets determined using Panther GOSlim Fisher’s overrepresentation test with false-discovery rate correction. The outer-most terms for each hierarchical GO family which was significantly enriched (P < 0.05) are shown, colored according to the cluster with which they were associated.

**Fig. S5:**
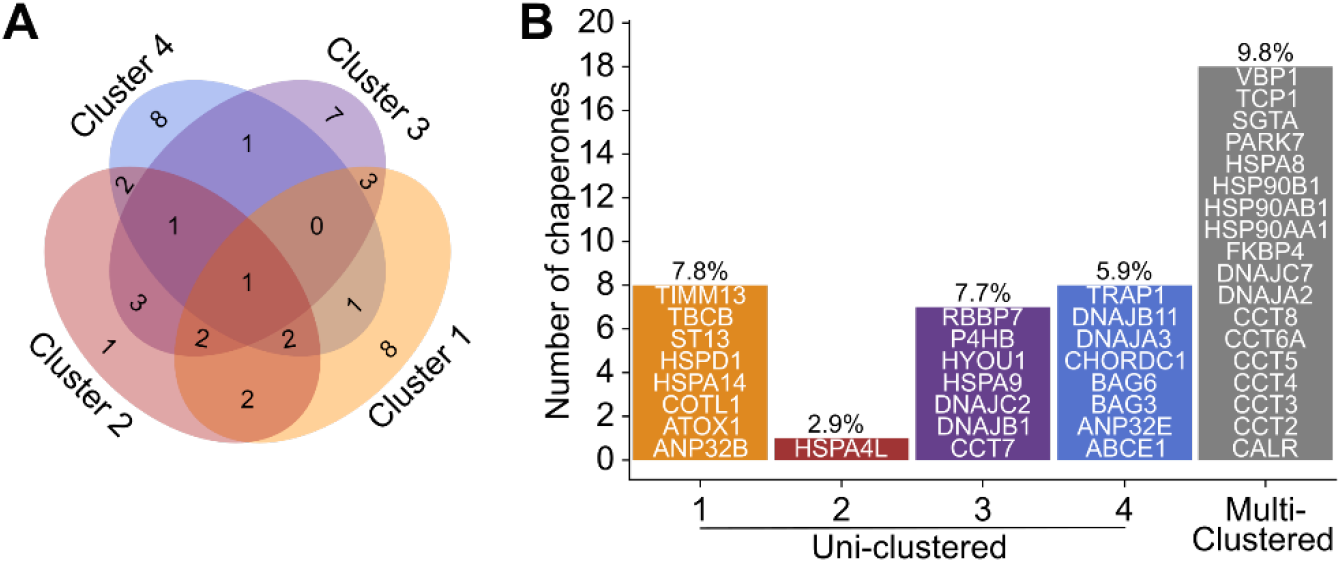
Enrichment of chaperone machinery among multi-clustered proteins. Relates to Fig. 4. (A) Venn diagram depicting proportion of chaperone proteins for which peptides were found in each cluster combination. (B) Number and proportion of proteins in each cluster associated with “chaperone-mediated protein folding” gene ontology term (GO:0061077). Gene names for individual proteins are listed within the bars.

**Fig. S6:**
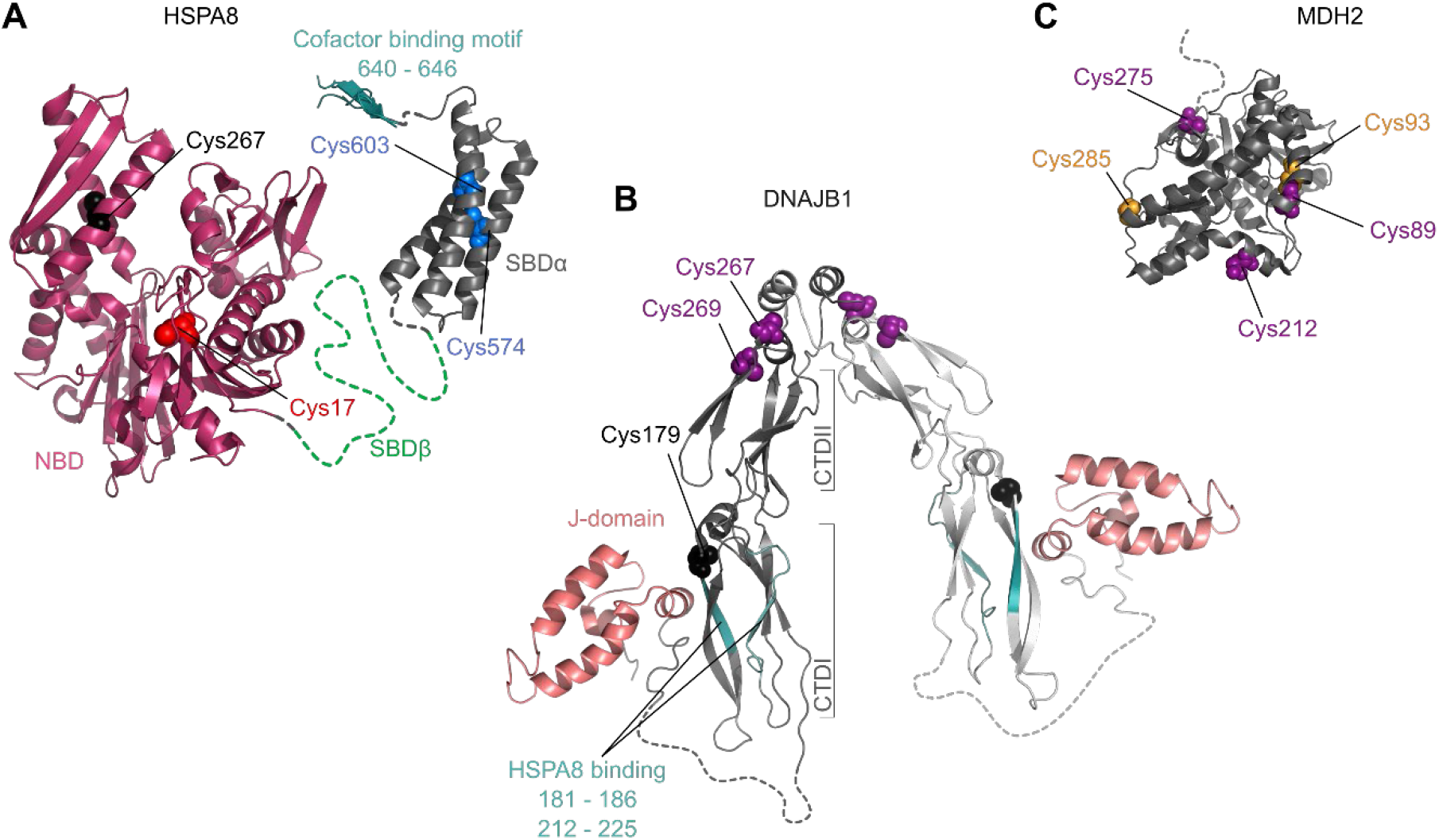
Structural models for chaperone client-binding reaction components. Relates to Fig. 4. (A) Ribbon structure of HSPA8 (models built with PDB 4H5R, 3AGY and 4KBQ). Nucleotide binding domain (NBD; ruby), substrate binding domains (SBDβ green, SBDα dark-grey) and cofactor binding motif (EEVD motif; teal) are shown on protein backbones. (B) Ribbon structure of DNAJB1 (model built with PDB 3AGZ and 1HDJ), with dimer composed of two monomers colored dark and light grey respectively. J-domain (salmon) and HSPA8 binding region (teal) are shown on protein backbones, and C-terminal domains (CTDI and CTDII) are indicated by brackets. (C) Ribbon structure of MDH2 (PDB 1MLD). In all panels, cysteine residues are labelled and colored according to the cluster their respective peptides were assigned (orange, red, purple and blue correspond to clusters 1 – 4 respectively, black was not observed), and dotted lines represent sequence regions with missing structural information.

**Table S1:**
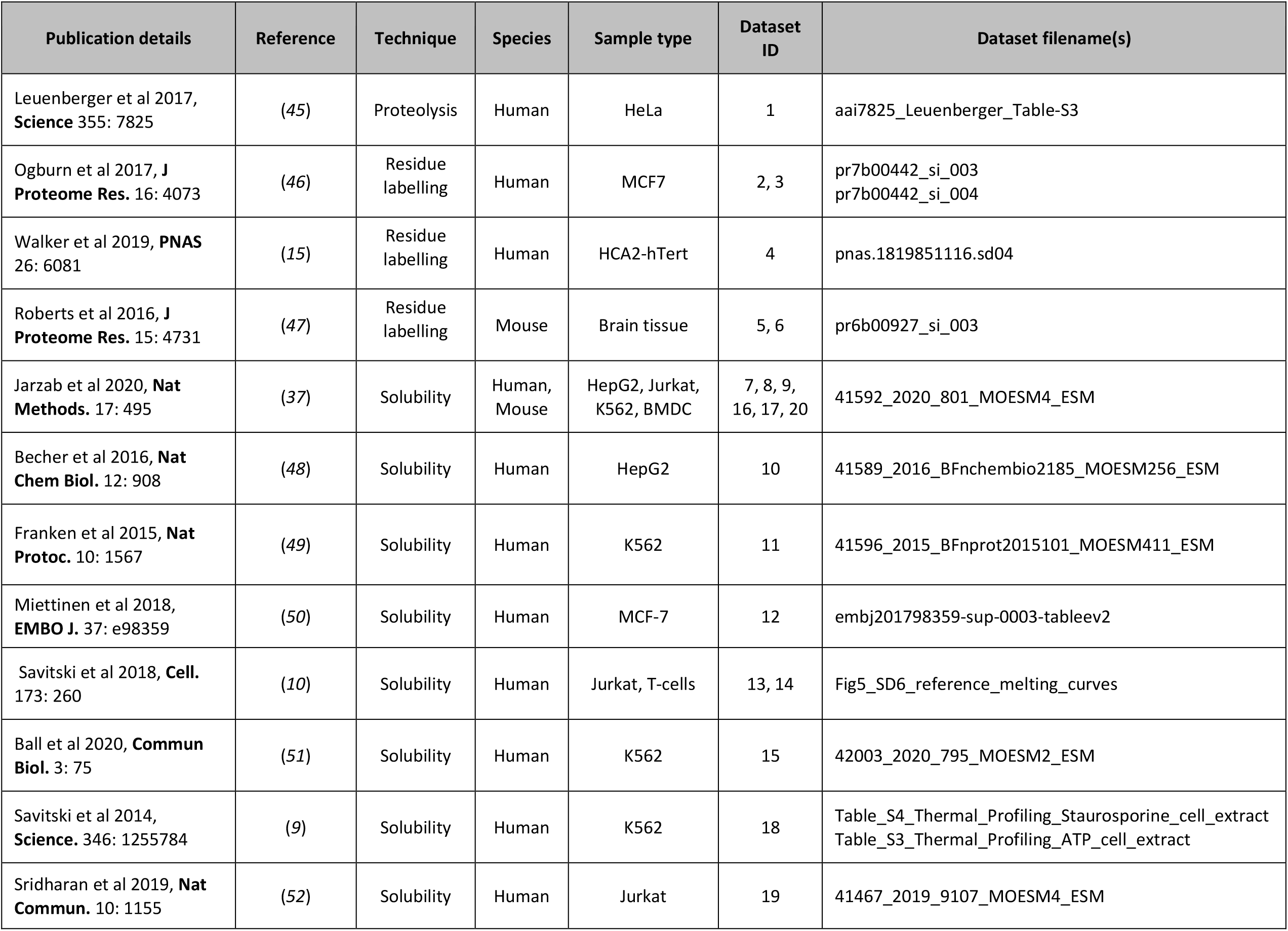
Published proteome stability dataset details.

## References and Notes

1. W. E. Balch, R. I. Morimoto, A. Dillin, J. W. Kelly, Adapting Proteostasis for Disease Intervention. Science. 319, 916–919 (2008).

2. G. G. Jayaraj, M. S. Hipp, F. U. Hartl, Functional Modules of the Proteostasis Network. Cold Spring Harb Perspect Biol. 12, a033951 (2020).

3. J. Labbadia, R. I. Morimoto, The Biology of Proteostasis in Aging and Disease. Annu Rev Biochem. 84, 435–464 (2015).

4. E. Braselmann, J. L. Chaney, P. L. Clark, Folding the proteome. Trends Biochem. Sci. 38, 337–344 (2013).

5. D. Cox, C. Raeburn, X. Sui, D. M. Hatters, Protein aggregation in cell biology: An aggregomics perspective of health and disease. Seminars in Cell & Developmental Biology (2018), doi:10.1016/J.SEMCDB.2018.05.003.

6. M. Z. Chen, N. S. Moily, J. L. Bridgford, R. J. Wood, M. Radwan, T. A. Smith, Z. Song, B. Z. Tang, L. Tilley, X. Xu, G. E. Reid, M. A. Pouladi, Y. Hong, D. M. Hatters, A thiol probe for measuring unfolded protein load and proteostasis in cells. Nat Commun. 8, 474 (2017).

7. F. Liu, H. Meng, M. C. Fitzgerald, Large-Scale Analysis of Breast Cancer-Related Conformational Changes in Proteins Using SILAC-SPROX. J. Proteome Res. 16, 3277–3286 (2017).

8. D. M. Molina, R. Jafari, M. Ignatushchenko, T. Seki, E. A. Larsson, C. Dan, L. Sreekumar, Y. Cao, P. Nordlund, Monitoring drug target engagement in cells and tissues using the cellular thermal shift assay. Science. 341, 84–87 (2013).

9. M. M. Savitski, F. B. M. Reinhard, H. Franken, T. Werner, M. F. Savitski, D. Eberhard, D. Martinez Molina, R. Jafari, R. B. Dovega, S. Klaeger, B. Kuster, P. Nordlund, M. Bantscheff, G. Drewes, Tracking cancer drugs in living cells by thermal profiling of the proteome. Science. 346, 1255784 (2014).

10. M. M. Savitski, N. Zinn, M. Faelth-Savitski, D. Poeckel, S. Gade, I. Becher, M. Muelbaier, A. J. Wagner, K. Strohmer, T. Werner, S. Melchert, M. Petretich, A. Rutkowska, J. Vappiani, H. Franken, M. Steidel, G. M. Sweetman, O. Gilan, E. Y. N. Lam, M. A. Dawson, R. K. Prinjha, P. Grandi, G. Bergamini, M. Bantscheff, Multiplexed proteome dynamics profiling reveals mechanisms controlling protein homeostasis. Cell. 173, 260–274.e25 (2018).

11. S. Schopper, A. Kahraman, P. Leuenberger, Y. Feng, I. Piazza, O. Müller, P. J. Boersema, P. Picotti, Measuring protein structural changes on a proteome-wide scale using limited proteolysis-coupled mass spectrometry. Nat Protoc. 12, 2391–2410 (2017).

12. C. S. H. Tan, K. D. Go, X. Bisteau, L. Dai, C. H. Yong, N. Prabhu, M. B. Ozturk, Y. T. Lim, L. Sreekumar, J. Lengqvist, V. Tergaonkar, P. Kaldis, R. M. Sobota, P. Nordlund, Thermal proximity coaggregation for system-wide profiling of protein complex dynamics in cells. Science. 359, 1170–1177 (2018).

13. I. Becher, A. Andrés-Pons, N. Romanov, F. Stein, M. Schramm, F. Baudin, D. Helm, N. Kurzawa, A. Mateus, M.-T. Mackmull, A. Typas, C. W. Müller, P. Bork, M. Beck, M. M. Savitski, Pervasive Protein Thermal Stability Variation during the Cell Cycle. Cell. 173, 1495–1507.e18 (2018).

14. T. J. Magliery, J. J. Lavinder, B. J. Sullivan, Protein stability by number: high-throughput and statistical approaches to one of protein science’s most difficult problems. Curr Opin Chem Biol. 15, 443–451 (2011).

15. E. J. Walker, J. Q. Bettinger, K. A. Welle, J. R. Hryhorenko, S. Ghaemmaghami, Global analysis of methionine oxidation provides a census of folding stabilities for the human proteome. PNAS. 116, 6081–6090 (2019).

16. S. M. Marino, V. N. Gladyshev, Cysteine function governs its conservation and degeneration and restricts its utilization on protein surfaces. J. Mol. Biol. 404, 902–916 (2010).

17. P. Busti, C. A. Gatti, N. J. Delorenzi, Some aspects of beta-lactoglobulin structural properties in solution studied by fluorescence quenching. Int. J. Biol. Macromol. 23, 143–148 (1998).

18. J. C. Bezdek, Pattern Recognition with Fuzzy Objective Function Algorithms (Springer Science & Business Media, 1981).

19. C. Döring, M.-J. Lesot, R. Kruse, Data analysis with fuzzy clustering methods. Computational Statistics & Data Analysis. 51, 192–214 (2006).

20. G. Chakafana, T. Zininga, A. Shonhai, The Link That Binds: The Linker of Hsp70 as a Helm of the Protein’s Function. Biomolecules. 9 (2019).

21. S. K. Sharma, P. De los Rios, P. Christen, A. Lustig, P. Goloubinoff, The kinetic parameters and energy cost of the Hsp70 chaperone as a polypeptide unfoldase. Nat Chem Biol. 6, 914–920 (2010).

22. R. Rosenzweig, N. B. Nillegoda, M. P. Mayer, B. Bukau, The Hsp70 chaperone network. Nat. Rev. Mol. Cell Biol. 20, 665–680 (2019).

23. N. B. Nillegoda, J. Kirstein, A. Szlachcic, M. Berynskyy, A. Stank, F. Stengel, K. Arnsburg, X. Gao, A. Scior, R. Aebersold, D. L. Guilbride, R. C. Wade, R. I. Morimoto, M. P. Mayer, B. Bukau, Crucial HSP70 co–chaperone complex unlocks metazoan protein disaggregation. Nature. 524, 247–251 (2015).

24. A. Ahmad, A. Bhattacharya, R. A. McDonald, M. Cordes, B. Ellington, E. B. Bertelsen, E. R. P. Zuiderweg, Heat shock protein 70 kDa chaperone/DnaJ cochaperone complex employs an unusual dynamic interface. Proc Natl Acad Sci U S A. 108, 18966–18971 (2011).

25. W. Han, P. Christen, Mechanism of the Targeting Action of DnaJ in the DnaK Molecular Chaperone System. J. Biol. Chem. 278, 19038–19043 (2003).

26. A. Mashaghi, S. Bezrukavnikov, D. P. Minde, A. S. Wentink, R. Kityk, B. Zachmann-Brand, M. P. Mayer, G. Kramer, B. Bukau, S. J. Tans, Alternative modes of client binding enable functional plasticity of Hsp70. Nature. 539, 448–451 (2016).

27. H. Suzuki, S. Noguchi, H. Arakawa, T. Tokida, M. Hashimoto, Y. Satow, Peptide-Binding Sites As Revealed by the Crystal Structures of the Human Hsp40 Hdj1 C-Terminal Domain in Complex with the Octapeptide from Human Hsp70. Biochemistry. 49, 8577–8584 (2010).

28. T. R. M. Barends, R. W. W. Brosi, A. Steinmetz, A. Scherer, E. Hartmann, J. Eschenbach, T. Lorenz, R. Seidel, R. L. Shoeman, S. Zimmermann, R. Bittl, I. Schlichting, J. Reinstein, Combining crystallography and EPR: crystal and solution structures of the multidomain cochaperone DnaJ. Acta Crystallogr D Biol Crystallogr. 69, 1540–1552 (2013).

29. J. C. Borges, H. Fischer, A. F. Craievich, C. H. I. Ramos, Low Resolution Structural Study of Two Human HSP40 Chaperones in Solution DJA1 from subfamily A and DJB4 from subfamily B have different quaternary structures. J. Biol. Chem. 280, 13671–13681 (2005).

30. R. Ch, O. Cl, F. Cy, T. Il, C. Dm, Conserved central domains control the quaternary structure of type I and type II Hsp40 molecular chaperones. J Mol Biol. 383, 155–166 (2008).

31. T. A. Määttä, M. Rettel, S. Sridharan, D. Helm, N. Kurzawa, F. Stein, M. M. Savitski, Aggregation and disaggregation features of the human proteome. Mol Syst Biol. 16 (2020).

32. X. Sui, D. E. V. Pires, A. R. Ormsby, D. Cox, S. Nie, G. Vecchi, M. Vendruscolo, D. B. Ascher, G. E. Reid, D. M. Hatters, Widespread remodeling of proteome solubility in response to different protein homeostasis stresses. Proc Natl Acad Sci USA. 117, 2422–2431 (2020).

33. E. W. J. Wallace, J. L. Kear-Scott, E. V. Pilipenko, M. H. Schwartz, P. R. Laskowski, A. E. Rojek, C. D. Katanski, J. A. Riback, M. F. Dion, A. M. Franks, E. M. Airoldi, T. Pan, B. A. Budnik, D. A. Drummond, Reversible, specific, active aggregates of endogenous proteins assemble upon heat stress. Cell. 162, 1286–1298 (2015).

34. A. Mateus, N. Kurzawa, I. Becher, S. Sridharan, D. Helm, F. Stein, A. Typas, M. M. Savitski, Thermal proteome profiling for interrogating protein interactions. Mol Syst Biol. 16 (2020) (available at https://www.ncbi.nlm.nih.gov/pmc/articles/PMC7057112/).

35. N. Kurzawa, I. Becher, S. Sridharan, H. Franken, A. Mateus, S. Anders, M. Bantscheff, W. Huber, M. M. Savitski, A computational method for detection of ligand-binding proteins from dose range thermal proteome profiles. Nature Communications. 11, 5783 (2020).

36. B. Seashore-Ludlow, H. Axelsson, T. Lundbäck, Perspective on CETSA Literature: Toward More Quantitative Data Interpretation. SLAS Discov. 25, 118–126 (2020).

37. A. Jarzab, N. Kurzawa, T. Hopf, M. Moerch, J. Zecha, N. Leijten, Y. Bian, E. Musiol, M. Maschberger, G. Stoehr, I. Becher, C. Daly, P. Samaras, J. Mergner, B. Spanier, A. Angelov, T. Werner, M. Bantscheff, M. Wilhelm, M. Klingenspor, S. Lemeer, W. Liebl, H. Hahne, M. M. Savitski, B. Kuster, Meltome atlas—thermal proteome stability across the tree of life. Nature Methods. 17, 495–503 (2020).

38. W. Kabsch, C. Sander, Dictionary of protein secondary structure: pattern recognition of hydrogen-bonded and geometrical features. Biopolymers. 22, 2577–2637 (1983).

39. W. G. Touw, C. Baakman, J. Black, T. A. H. te Beek, E. Krieger, R. P. Joosten, G. Vriend, A series of PDB-related databanks for everyday needs. Nucleic Acids Res. 43, D364–D368 (2015).

40. B. Mészáros, G. Erdos, Z. Dosztányi, IUPred2A: context-dependent prediction of protein disorder as a function of redox state and protein binding. Nucleic Acids Res. 46, W329–W337 (2018).

41. D. Szklarczyk, A. L. Gable, D. Lyon, A. Junge, S. Wyder, J. Huerta-Cepas, M. Simonovic, N. T. Doncheva, J. H. Morris, P. Bork, L. J. Jensen, C. von Mering, STRING v11: protein-protein association networks with increased coverage, supporting functional discovery in genome-wide experimental datasets. Nucleic Acids Res. 47, D607–D613 (2019).

42. Z. Chen, P. Zhao, F. Li, A. Leier, T. T. Marquez-Lago, Y. Wang, G. I. Webb, A. I. Smith, R. J. Daly, K.-C. Chou, J. Song, iFeature: a Python package and web server for features extraction and selection from protein and peptide sequences. Bioinformatics. 34, 2499–2502 (2018).

43. P. Virtanen, R. Gommers, T. E. Oliphant, M. Haberland, T. Reddy, D. Cournapeau, E. Burovski, P. Peterson, W. Weckesser, J. Bright, S. J. van der Walt, M. Brett, J. Wilson, K. J. Millman, N. Mayorov, R. J. Nelson, E. Jones, R. Kern, E. Larson, C. J. Carey, I. Polat, Y. Feng, E. W. Moore, J. VanderPlas, D. Laxalde, J. Perktold, R. Cimrman, I. Henriksen, E. A. Quintero, C. R. Harris, A. M. Archibald, A. H. Ribeiro, F. Pedregosa, P. van Mulbregt, SciPy 1.0: fundamental algorithms for scientific computing in Python. Nature Methods. 17, 261–272 (2020).

44. Y. Perez-Riverol, A. Csordas, J. Bai, M. Bernal-Llinares, S. Hewapathirana, D. J. Kundu, A. Inuganti, J. Griss, G. Mayer, M. Eisenacher, E. Pérez, J. Uszkoreit, J. Pfeuffer, T. Sachsenberg, S. Yilmaz, S. Tiwary, J. Cox, E. Audain, M. Walzer, A. F. Jarnuczak, T. Ternent, A. Brazma, J. A. Vizcaíno, The PRIDE database and related tools and resources in 2019: improving support for quantification data. Nucleic Acids Res. 47, D442–D450 (2019).

45. P. Leuenberger, S. Ganscha, A. Kahraman, V. Cappelletti, P. J. Boersema, C. von Mering, M. Claassen, P. Picotti, Cell-wide analysis of protein thermal unfolding reveals determinants of thermostability. Science. 355 (2017).

46. R. N. Ogburn, L. Jin, H. Meng, M. C. Fitzgerald, Discovery of Tamoxifen and N-Desmethyl Tamoxifen Protein Targets in MCF-7 Cells Using Large-Scale Protein Folding and Stability Measurements. J. Proteome Res. 16, 4073–4085 (2017).

47. J. H. Roberts, F. Liu, J. M. Karnuta, M. C. Fitzgerald, Discovery of age-related protein folding stability differences in the mouse brain proteome graphical abstract HHS public access. J Proteome Res. 15, 4731–4741 (2016).

48. I. Becher, T. Werner, C. Doce, E. A. Zaal, I. Tögel, C. A. Khan, A. Rueger, M. Muelbaier, E. Salzer, C. R. Berkers, P. F. Fitzpatrick, M. Bantscheff, M. M. Savitski, Thermal profiling reveals phenylalanine hydroxylase as an off-target of panobinostat. Nature Chemical Biology. 12, 908–910 (2016).

49. H. Franken, T. Mathieson, D. Childs, G. M. A. Sweetman, T. Werner, I. Tögel, C. Doce, S. Gade, M. Bantscheff, G. Drewes, F. B. M. Reinhard, W. Huber, M. M. Savitski, Thermal proteome profiling for unbiased identification of direct and indirect drug targets using multiplexed quantitative mass spectrometry. Nature Protocols. 10, 1567–1593 (2015).

50. T. P. Miettinen, J. Peltier, A. Härtlova, M. Gierliński, V. M. Jansen, M. Trost, M. Björklund, Thermal proteome profiling of breast cancer cells reveals proteasomal activation by CDK4/6 inhibitor palbociclib. The EMBO journal, e98359 (2018).

51. K. A. Ball, K. J. Webb, S. J. Coleman, K. A. Cozzolino, J. Jacobsen, K. R. Jones, M. H. B. Stowell, W. M. Old, An isothermal shift assay for proteome scale drug-target identification. Communications Biology. 3, 1–10 (2020).

52. S. Sridharan, N. Kurzawa, T. Werner, I. Günthner, D. Helm, W. Huber, M. Bantscheff, M. M. Savitski, Proteome-wide solubility and thermal stability profiling reveals distinct regulatory roles for ATP. Nature Communications. 10, 1155 (2019).

